# Enhanced survival and lower growth in assisted gene flow corals during a marine heatwave driven by local adaptation

**DOI:** 10.64898/2026.01.13.698983

**Authors:** Alex Macadam, Ira R. Cooke, Patrick W. Laffy, Jan M. Strugnell, Carys Morgans, Andrea Severati, Kate M. Quigley

## Abstract

Recent increases in the frequency of mass coral bleaching and mortality events have caused shifts in species composition and rapid global reef decline. Genetic-based techniques, like assisted gene flow (AGF), offer the potential to enhance corals’ climate readiness by introducing genetic variation associated with heat tolerance into vulnerable populations. This method remains untested in the wild during a marine heatwave. To this end, we selectively bred AGF corals by crossing parent *Acropora tersa* corals collected across five Great Barrier Reef (GBR) locations spanning a thermal and bleaching gradient. Breeding produced intra- and inter-regional offspring that were then deployed to a central GBR reef. This deployment reef was subsequently hit by a marine heatwave (>31.4°C for 17 days and >6 Degree Heating Weeks). Heat tolerance-associated traits (survival, growth, and bleaching) were compared between AGF offspring and a native control cross over the 95 days in the wild. Overall, AGF offspring had 4× greater odds of survival compared to controls, and 2× greater odds compared to northern juveniles. Growth rates were 14% higher in control juveniles compared to the mean growth rate across all AGF crosses. Interestingly, heat-stress assays (32°C) of the parental adult corals showed similar patterns to their offspring, with higher survival and lower bleaching of corals from warmer northern reefs. Taken together, these findings providing critical support that the selective breeding of AGF corals can provide enhanced survival under marine heatwave conditions in the wild. It also suggests further evaluation is needed for trait-specific responses to effectively match donor and recipient reefs in restoration planning.

## INTRODUCTION

Coral reefs globally are under increasing threat from climate change and other anthropogenic stressors (De’Ath et al., 2012; Eddy et al., 2021), in which rising ocean temperatures pose the greatest threat to their survival (Hughes et al., 2018c; Van Woesik et al., 2022). On the Great Barrier Reef (GBR), recurrent mass bleaching events driven by accumulated heat stress and subsequent breakdown of the coral-algal symbiosis (Douglas, 2003) have become the primary cause of degradation (Donner et al., 2017; Hughes et al., 2018a; Skirving et al., 2019). These events not only reduce coral cover but are also fundamentally reshaping reef assemblages (Hughes et al., 2018b; Stuart-Smith et al., 2018).

There is evidence that rapid changes to physiology (i.e., phenotypic plasticity) can support short-term resistance (Drury et al., 2022c), many coral species are likely already close to their thermal limits as oceans continue to warm (Arias, 2021). Along with the coral microbiome (Bourne et al., 2016; Quigley et al., 2020b), coral host genetics also plays a key role in modulating the thermal tolerance of the organism (Dixon et al., 2015; Macadam et al., 2025). For example, substantial variation in heat tolerance exists across reefs on the GBR in at least five coral species (Denis et al., 2024; Jurriaans & Hoogenboom, 2019; Naugle et al., 2024; Ulstrup et al., 2006). There is evidence of adaptive potential in some coral populations (Matz et al., 2018), however, the speed at which coral may adapt to increasingly hot seawater temperatures may not be sufficient to match the pace of ocean warming (Quigley et al., 2019). The continued degradation of reefs has spurred interest and development in genetic-based conservation and restoration strategies to bolster coral resilience (McLeod et al., 2019; Shaver et al., 2022).

Several methodologies exist to restore corals, including coral fragmentation and propagation (Boström-Einarsson et al., 2020; Howlett et al., 2023). Although these approaches can boost coral cover and biomass, they may not necessarily increase heat tolerance (Boström-Einarsson et al., 2020). As such, out-planted corals may remain vulnerable to marine heatwaves, limiting gains in overall reef resilience (Mulà et al., 2025; Szereday et al., 2025). To enhance stress tolerance, particularly to heat, genetic-based methods have been proposed to either increase heat tolerance at local reefs or facilitate the spread of heat tolerance via the transfer of adaptive variation (Baums et al., 2022). One such intervention, known as assisted gene flow (AGF), is defined as the intentional translocation of individuals within a species range to facilitate adaptation to anticipated future conditions at that location (Aitken & Whitlock, 2013). The potential for AGF as a conservation strategy on coral reefs of the GBR has been demonstrated through laboratory studies on larval and juvenile corals whose parents have been sourced from regions with different thermal histories (Dixon et al., 2015; Howells et al., 2021; Macadam et al., 2025; Quigley et al., 2020b; van Oppen & Quigley, 2022; Weeriyanun et al., 2022). Initial tests of AGF juveniles in the wild suggested that these benefits could be transferred from the lab to the field (Quigley et al., 2021). A crucial test, however, is whether corals produced using AGF methods exhibit higher heat tolerance or generally higher fitness during a marine heatwave in the wild. This information is crucial to determine whether enhanced survival observed in laboratory experiments stands up during a marine heatwave outside of the lab.

Here, we generated empirical data on AGF produced and native coral offspring across key fitness traits (survival, growth, and bleaching) collected during a marine heatwave. We found that overall, coral offspring bred using AGF methods showed enhanced survival relative to native, “control” corals under the heat stress conditions of the marine heatwave. However, the enhanced survival came at a cost of decreased growth. These results demonstrate the feasibility of AGF in the field whilst also highlighting that trade-offs in key fitness traits may occur. This study provides critical insights for practitioners employing conservation strategies aimed at balancing gains in heat tolerance with potential costs, like reduced growth rates.

## MATERIALS AND METHODS

### Coral collection

Gravid colonies of *Acropora tersa* (Rassmussen et al., 2025) were collected from reefs along a latitudinal gradient on the GBR. These reefs were as follows: Wood (northern, outer shelf; 11.811805⁰S, 143.981617⁰E), Martin (northern, mid-shelf; 14.77130⁰S, 145.36940⁰E), Arlington (northern, mid-shelf; 16.64293⁰S, 146.10841⁰E), Eumilli Island (Fantome Island) in the Palm Island group (central, inshore-mid shelf; 18.65907⁰S, 146.50455⁰E) and Davies (central, mid – outer shelf; −18.81978⁰S, 147.64563⁰E). Colonies were collected over two weeks in November 2021 (Fig. 1A). Reefs were chosen for their variation in annual mean temperatures (Fig. 1B). Coral colonies were transported to the National Sea Simulator (SeaSim) at the Australian Institute of Marine Science, Townsville, Australia, for spawning. Coral colonies from each reef were maintained in separate 1,000 L holding tanks within a semi-open recirculating system. Water temperature was regulated using a heat exchanger set to the mean historical temperature for each collection reef, based on site-specific data extracted from eReefs (https://portal.ereefs.info/map). The tanks were supplied with flow-through water at a rate of 6 L/min. Positioned outdoors, colonies received natural daylight, with shade cloth reducing the light intensity to ∼155 PAR at midday.

**Figure 1:**
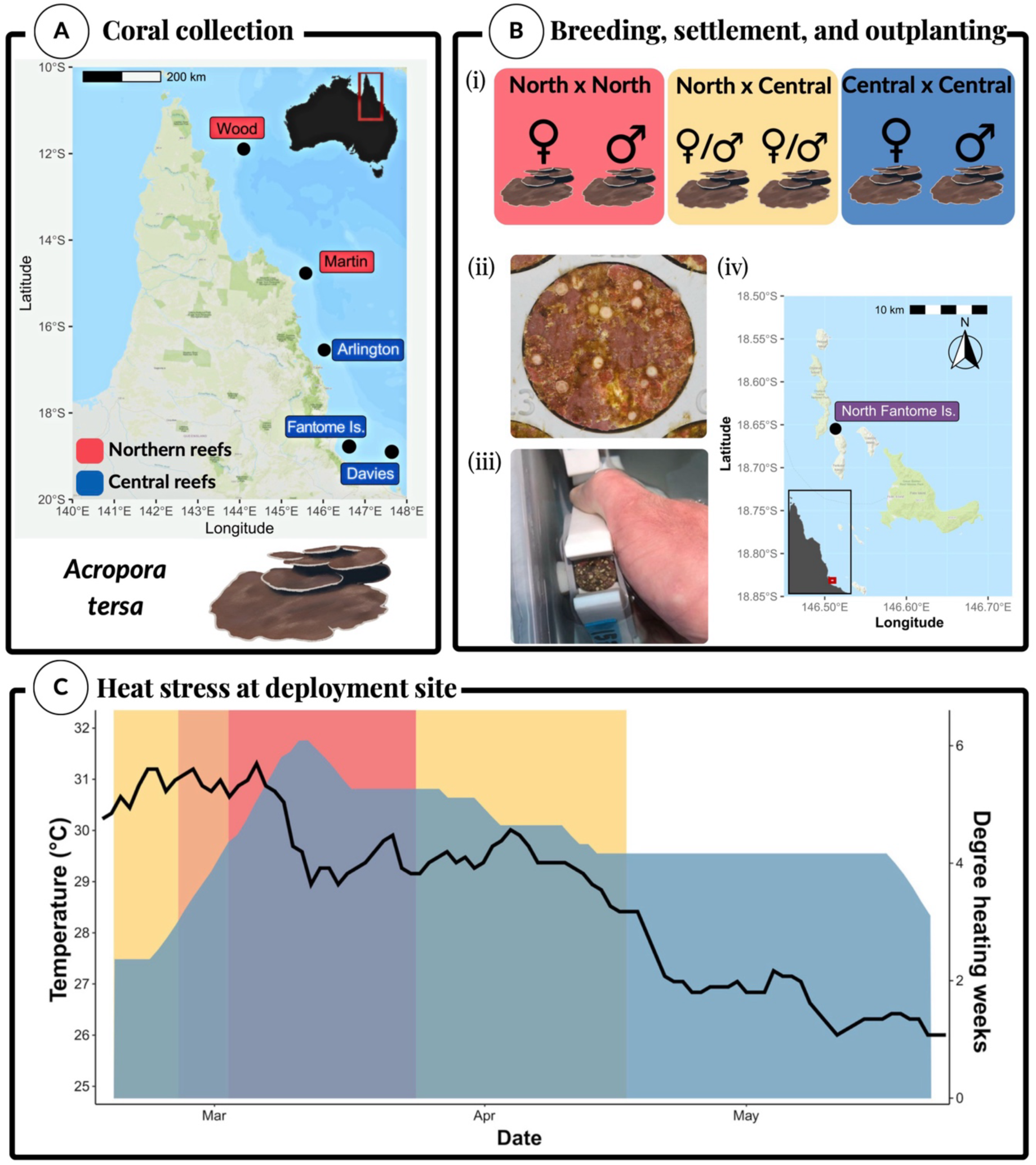
Location of parental corals, breeding design, and heat stress experienced during deployment. (A) Map of the Great Barrier Reef, Australia, showing the *Acropora tersa* coral collection sites. Adult corals were collected from two northern sites (red) and three central sites (blue) on the Great Barrier Reef. (B) Diagram of coral breeding, settlement, and deployment. (i) Gametes were mixed to produce intra-region North × North & Central × Central crosses, and inter-region (North × Central and Central × North) crosses. Larvae were settled onto aragonite plugs (ii), which were fitted into specially designed devices (iii). Twenty-eight-day old juveniles were put on tables that were fixed to the seafloor at the northern tip of Fantome Island, the Palm Island Group, the Great Barrier Reef, Australia (iv) on 19th of February 2022. (C) The heat stress experienced by the coral juvenile during deployment. Temperature (black line), degree heating weeks (DHW; blue bars), and NOAA Coral Reef Watch Bleaching Alert level (background; yellow = ‘Watch’, orange = ‘Warning’, red = ‘Alert Level 1’). Juveniles were retrieved from the field after 95 days.

### Coral breeding and larval rearing

Coral spawning times were estimated using the Indo-Pacific coral spawning database (Baird et al., 2021). Spawning, fertilisation and rearing followed established methods (Quigley et al., 2016). Briefly, once corals were visibly set (when egg and sperm bundles become visible in the polyp mouth), each colony was isolated in a separate container to reduce the chance of cross contamination. Gamete bundles were collected from the water surface and transferred into a 120µm filter. The bundles were washed three times using 0.2 µm filtered sea water (FSW) to separate the eggs and the sperm. Sperm concentration was estimated using an automated sperm counter (Computer - Assisted Semen Analysis-CASA equipment), and sperm concentrations were adjusted so that sperm was at a concentration of 1 x 10^6^. To produce corals with both parents from a central reef (hereby referred to as intra-region central juveniles) and northern reefs (hereby referred to as intra-region northern juveniles), sperm and eggs from six colonies were mixed per region cross. To produce corals with parents from different reefs (hereby referred to as inter-region juveniles), corals from different regions were required to spawn on the same night, as gametes could not be stored for longer than a few hours without producing confounding effects. Due to differences in spawning times, some crosses could not be produced. This included crossing Fantome Island corals with corals from other reefs.

Seven reproductive crosses of coral offspring groups were produced. These include offspring produced from crossing central dams × central sires (intra-region central juveniles). These are listed as the following: Fantome × Fantome (FI×FI), Davies × Davies (DA×DA), and Davies × Arlington (DA×AR). FI×FI also acted as native controls at our deployment site at Fantome. Juveniles were produced by crossing northern dams × northern sires (intra-region northern juveniles). These were: Martin × Martin (MA×MA), and Wood × Martin (WD×MA). Finally, AGF offspring were produced, made up of crossing northern and central dams and sires (inter-region juveniles). These were as follows: Martin × Davies (MA×DA) and Davies × Martin (DA×MA).

Fertilisation success was confirmed two hours after adding the sperm by checking embryos under a microscope. Embryos for each cross were transferred into rearing tanks with flow through filtered sea water (FSW). After 24 h, each tank received constant flow-through of 0.2 μm FSW and an air bubbler to reduce water stagnation. Larvae were reared in cones until they were competent to settle, which was after 10 days (Pollock et al., 2017).

### Coral juvenile settlement and deployment

Aragonite plugs and terracotta tiles were prepared as a settlement substrate for coral larvae by conditioning them for six weeks in tanks containing crustose coralline algae, a known coral settlement inducer (Heyward & Negri, 1999). Plugs were fixed into specially designed holding racks. The tiles had holes drilled through the centre and were fixed onto metal rods with 2 cm separators between each. Plug racks and tile rods were placed into 50 L tanks with flow-through filtered sea water set at 27.5°C. Each larval cross was added to a specific tank, in which larvae could settle onto the aragonite plugs and tiles. Plugs and tiles were maintained in these conditions for 28 days until deployment. Two days prior to deployment, the aragonite plugs were fitted into devices specifically designed to enhance the survival of deployed coral juveniles by minimizing predation (Whitman et al., 2024). These devices were then fixed to specially designed holding racks (Fig. 1B), each labelled with unique identifiers. The temperature in the holding tanks with the tiles and devices were then increased over time (ramped by 0.5°C per day) to the current temperature of the deployment site (30.0°C).

At the day of deployment, juvenile corals were 28-days old. These juveniles were deployed to north Fantome Island, the Palms Islands, Australia, on the 19^th^ of February 2022. The holding racks of devices with coral juveniles were deployed to the reef using SCUBA (Fig. 1C), following Quigley et al. (2021). The racks of devices were placed onto frames that were secured onto the sand using metal rods. Each frame (1 x 2.9 m) held 12 racks, each rack held 12 devices, with each device holding three plugs. In total n = 864 plugs were deployed, which consisted of n = 7,750 juvenile corals.

### Coral juvenile survival, colour, and growth

Three key fitness traits – survival, growth, and colour – were measured in the coral juveniles. These traits were measured before deployment and after 95 days in the field. Prior to deployment, juveniles were photographed using a Nikon D810 camera body, Nikon AFS 60 mm micro lens, with each photo including a 2 cm scale bar and CoralWatch Health Chart (Siebeck et al., 2006) as described in Quigley et al. (2021). On day 95, devices were retrieved using SCUBA, and plugs were removed from the devices and placed into a plug tray for photographs. In the field, plugs were photographed with an Olympus Tough TG-6 camera with an Olympus PT-058 waterproof housing. A CoralWatch Health chart was photographed with the juveniles to use as a colour reference. The camera was fixed onto a PVC frame mounted onto a flow-through water tray for constant angle and distance from camera to subject. Consistent camera and flash settings were also used. Survival was measured by comparing the number of juveniles at deployment and number of juveniles alive by the end of deployment period. Growth was measured as the percent difference in size from the start and end of the deployment period using the program Fiji (Rueden & Eliceiri, 2019). Coral bleaching was also measured using Fiji. Coral juveniles were selected on images set to red, green, blue (RGB) scale. Mean grey value of the selected juveniles was calculated, where 0 is black and 255 is white (DeMerlis et al., 2022; Wang et al., 2022). This value was then corrected in comparison to a reference to account for differences between images and then converted to a scale of 0 to 100, where 0 was equal to white and 100 was equal to black.

### Field and historic environmental data

During the deployment, two Hobo Pendant 64K Temp-Light Data Loggers were attached to each the outplant tables to record temperature and light intensity over time. NOAA Coral Reef Watch data was also used to assess Degree Heating Weeks (DHW) and Bleaching Alert Areas (BAAs) experienced at the site (see NOAA Coral Reef Watch, 2000). Briefly, the Bleaching Alert statuses are defined as follows: No Stress (no heat stress or bleaching is present), Bleaching Watch (low-level heat stress), Bleaching Warning (possible bleaching), Bleaching Alert Level 1 (significant bleaching likely) and Bleaching Alert Level 2 (severe bleaching and significant mortality likely). Historic temperature data was extracted from eReefs (https://portal.ereefs.info/map) for each collection site (CSIRO, 2020).

### Adult heat stress experiment

After spawning, the adult colonies of *A. tersa* were also tested for their thermal tolerance. Colonies from each reef were held at the SeaSim facility in aquarium systems synchronized to each reef’s historic temperature regime (data extracted from eReefs https://portal.ereefs.info/map for each collection site; CSIRO, 2020). A band saw with a diamond encrusted blade was used to cut similar sized coral fragments (∼5 cm). Fragments were then superglued onto aragonite plugs to provide at least six replicates for each genotype per temperature. Coral fragments were then allowed to acclimate for two weeks to allow time for the healing of fragmenting wounds. After this, corals were randomly distributed throughout replicate 50 L acrylic aquaria such that each genotype was represented at least once in each tank. Each aquarium operated as an independent flow-through system supplied with ultrafiltered, temperature-controlled natural seawater at 0.8 L/min (approximately one full turnover per hour). Aquaria were completely isolated from one another and were immersed in individual water jackets to stabilise temperature conditions, with internal pumps providing continuous water circulation. Lighting for each aquarium was set to a 24-hour light/dark cycle (sunrise: 06:00 and sunset: 16:00), 2-hours sunrise/sunset ramp time with photosynthetically active radiation (PAR) measuring at ∼150 PAR. The growth of fouling algae was cleaned every second day to support the health of corals. Feeding of corals with *Artemia nauplii* and algae was carried out at 15:00 each day.

A temperature of 32°C was chosen for the heat stress treatment (2°C above the Maximum Monthly Mean (MMM) of the warmest reef, Wood reef = 29.99°C), to ensure heat-induced bleaching occurred. The temperature in the hot treatment was increased by 0.5°C/day from ambient (27.5°C) to 32°C over a period of nine days for all aquaria (n = 6 tanks). The temperature of the ambient treatment aquaria remained at 27.5°C (n = 6 tanks). In summary, the resulting number of tanks were as follows: n = 6 ambient temperature, n = 6 hot temperature. The first day of measurements were taken the day after the hot aquaria reached the treatment temperature of 32°C.

Survival and colour (bleaching) was assessed using an Olympus TG5 camera fixed to a tripod set to a fixed position to ensure consistent positioning and lighting conditions. Images were taken alongside a CoralWatch Health Monitoring Chart (Siebeck et al., 2006) under standardized settings (ISO: 125, f/2.0, shutter speed: 1/25, no flash). The coral fragment colour was matched to the closest category on the CoralWatch “D” coral health chart by the same observer to minimise bias. The end of the experiment was set at when the first coral genotype reached a mean value of 50% mortality across the treatment tanks, which occurred after seven days.

### Statistical analysis

Statistical analyses of survival were performed using hierarchical logistic models in a Bayesian framework within the brms package (Bürkner, 2017) with ‘RStan’ (Guo et al., 2015). Separate models were fit for adults, larvae, and juveniles. The effect of temperature and reef identity on adult survival was analysed by fitting adult survival to a binomial distribution (number of live and dead fragments at each timepoint), where reef and temperature were included as fixed effects along with terms to account for their interaction. Tank was included as a random effect. Survival of coral juveniles was analysed by fitting juvenile survival to a binomial distribution (number of live and dead juveniles at each timepoint), where reef cross was included as a fixed effect, and rack (the device holding the plugs) was included as a random effect.

Change in colour (bleaching) and size were calculated for each individual juvenile. Growth and colour change were analysed using a Generalised linear mixed model (GLMM) in a Bayesian framework for each trait with Gaussian distribution using R package ‘brms’ (Bürkner, 2017) with reef cross as a fixed effect and rack as a random effect.

All models included three No-U-Turn (Markov chain Monte Carlo; MCMC) chains of 5000 iterations, thinned to a rate of 10, with a burn-in period of 2500. MCMC mixture and convergence were assessed using trace plots, autocorrelation plots, Rhat and effective sample size diagnostics. Model assumptions were assessed using residual diagnostics using the package ‘DHARMa’ (Hartig, 2017). All model diagnostics passed, and values derived from histograms indicated that all Rhat parameters ≤1.008, suggest strong convergence. Models were iteratively fitted using leave-one-out (LOO) cross-validation to assess fit. Bayesian exceedance probabilities (P_e_) are reported, which refers to the percent (%) confidence that the model has a greater posterior probability than any other model tested (Stephan et al., 2009). Credibility intervals are plotted for each treatment. The width of confidence intervals for the exceedance probability convey the stability of a result (Segal, 2021). The P_e_ scale adopted for this paper is: >0.85 = weak evidence, >0.9 = evidence, >0.975 = strong evidence. All results were visualised using the ‘ggplot2’ package (Wickham et al., 2016) in R (version 4.2.1; R Core Team, 2022) and all code used to execute the statistical analyses is available online (github.com/alexmacadam241/AGF-Outplanting).

## RESULTS

### Historical and contemporary temperature trends at collection and outplant reefs

Collections of adult *Acropora tersa* spanned a latitudinal gradient from the far northern sector of the GBR (11.8°S Wood Reef) to the central sector (18.8°S Davies Reef; Fig. 1A). The northern GBR sampling sites consisted of Wood and Martin reefs. The central GBR sites were Arlington and Davies reef, and Fantome Island. Overall, the key temperature metrics differed significantly amongst these sites, including mean Sea Surface Temperatures (SST), SST anomalies, DHW and BAA (Kruskal-Wallis: all *p <* 2.2×10⁻¹⁶; Supplementary Fig. S1). Pairwise comparisons revealed consistent latitudinal patterns. For example, Fantome and Davies showed no significant differences in SST (post-hoc Dunn test, Bonferroni *p*-adjusted: Z = 0.82, p = 1.0), SST anomalies (Z = 2.42, p = 0.15), DHW (Z = −2.33, *p =* 0.20), or BAA (Z = −0.11, p = 1.00), but both reefs reported significantly lower values for all metrics compared to Wood and Martin (*p* < 0.001). This was true, with the exception of SST anomalies, where Fantome had significantly fewer anomalies compared to Martin (Z = −3.22, *p* = 0.01), but not Davies (Z = −0.45, *p* = 1.00; Supplementary Fig. S1).

After juvenile deployment, a marine heatwave occurred at the outplant site (Fantome, Fig. 1A), thus creating a natural heat stress experiment. At the start of the experiment (the 19^th^ of February 2022), the deployment site experienced a daily mean SST of 29.53°C ± 0.04, exceeding the historic MMM that had been set for 18 of the past 20 years (Supplementary Table 1 & 2). Only the MMMs recorded in February of 2016 and 2017 were higher – which corresponded to years in which the GBR experienced mass bleaching and mortality events (Hughes et al., 2018c). Over the 91-day deployment period, SST ranged from 26.13°C ± 0.02 (the 24^th^ of May 2022) to a peak of 31.37°C ± 0.07 (the 9^th^ March 2022; Fig. 1C). By the 9^th^ of March, temperatures surpassed the historic MMM for Fantome (30.49°C ± 0.08; years 2000 - 2021). Additionally, temperatures also surpassed all yearly MMMs at all adult collection sites (Supplementary Table 1). At Fantome, SST exceeded this historic MMM for more than 20 days, equivalent to NOAA Bleaching Alert Level 1 and a cumulative total of six DHW (Fig. 1C).

### Fitness measurements of coral juveniles during a marine heatwave

#### a. Survival

By the end of the 91 days of deployment and the marine heatwave, intra-region northern juveniles had 1.9× greater odds of survival compared to intra-region central juveniles (Fig. 2A; P_e_ = 1.00, Bayesian Generalised Linear Mixed Model; BGLMM). Furthermore, inter-region juveniles had 4× greater odds of survival than the control intra-region central juveniles (P_e_ = 1.00, BGLMM), and 2× greater odds of survival than intra-region northern corals (P_e_ = 0.997, BGLMM). Juveniles from the native control cross, FI×FI, had 11% higher odds of survival compared to DA×DA juveniles (P_e_ = 1.00, BGLMM), the only other intra-region central cross.

**Figure 2:**
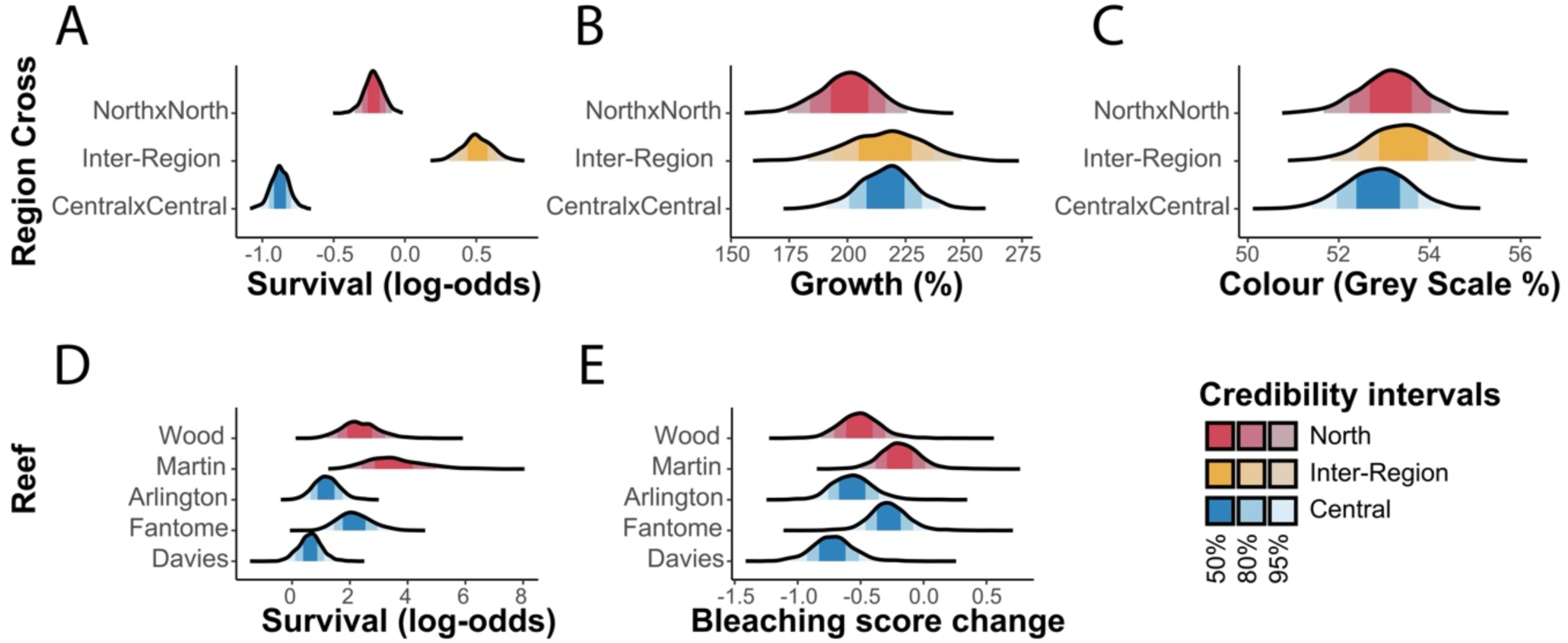
Survival and response to heat stress of *Acropora tersa* juvenile and adult corals. Distribution of estimated marginal means (EMM) for: (A) survival (log-odds), (B) growth (%), and (C) final colour (grey scale as a proxy for bleaching score) of juvenile corals at the end of deployment (95 days). (D) Survival (log-odds) and (E) bleaching (change in Coral Health Chart colour score D scale) of adult corals in the hot (32°C) treatment. Higher grey scale percent means more pigmentation. Colours represent the region; red = north, blue = central, and yellow = inter-region crosses. Intervals represent EMM 50%, 80% and 95% credibility intervals.

During this period of the marine heatwave, the mean survival was lowest in the control intra-region central juveniles (mean 29.6% ± 0.7 SE). Specifically, DA×DA had the lowest survival (27.5% ± 0.9) followed by FI×FI (29.0% ± 1.1; Supplementary Fig. S2A). Mean survival was highest in inter-region juveniles (63.0% ± 2.1). Specifically, DA×MA had the highest survival (78.2% ± 3.8) followed by DA×AR (69.3% ± 3.7) and MA×DA (58.6% ± 2.4). Mean survival of intra-region northern juveniles was lower than inter-region juveniles, but higher than intra-region central juveniles (northern = 44.4% ± 0.9). Specifically, WD×MA juveniles had a mean survival of 59.5% ± 2.5 and MA×MA had a mean survival of 42.1% ± 1.0.

#### b. Growth rates

Growth rates over 91 days of deployment increased by a mean of 209.3% ± 3.9. On average, intra-region northern juveniles grew 16% less compared to intra-region central juveniles (Fig. 2B; P_e_ = 0.97, BGLMM). The same intra-region northern juveniles also grew 16% less compared to inter-region juveniles (Fig. 2B; P_e_ = 0.87, BGLMM). Growth was not significantly different between inter-region and control intra-region central juveniles (P_e_ = 0.51, BGLMM).

At the regional scale, the mean growth rate was lowest in intra-region northern juveniles (northern = 198.0% ± 5.6). Specifically, MA×MA had the lowest growth rate (190.5% ± 6.1) followed by WD×MA (234.7% ± 14.5; Supplementary Fig. S2B). Mean growth rate was highest in inter-region juveniles (inter-region = 218.4% ± 12.0). Specifically, MA×DA had the highest growth rate (224.2% ± 14.8) followed by DA×MA (202.2% ± 18.4). Mean growth rate of intra-region central juveniles was lower than inter-region juveniles, but higher than intra-region northern juveniles (central = 217.8% ± 6.0). Specifically, the native juveniles at the outplant site, FI×FI, had the highest average growth rate (248.5% ± 10.1), followed by DA×AR (207.8% ± 16.2), and DA×DA (195.9% ± 8.0). These control FI×FI juveniles had a 46% higher growth rate compared to the other central intra-region cross DA×DA (P_e_ = 1.00, BGLMM).

#### c. Bleaching

Over the course of the deployment, juveniles became darker (measured as percentage of mean grey scale = 53.5% ± 0.2). Colour did not differ between northern and central intra-region juveniles (Fig. 2C; P_e_ = 0.82, BGLMM) or between northern intra-region and inter-region juveniles (P_e_ = 0.73, BGLMM). Conversely, intra-region central juveniles were lighter compared to inter-region juveniles (Fig. 2C; P_e_ = 0.88, BGLMM), indicating that intra-region central juveniles were more bleached compared to inter-region juveniles.

Intra-region central juveniles were the lightest in colouration (53.1% ± 0.2). Specifically, DA×DA were the lightest (52.7% ± 0.3; Supplementary Fig. S2C), followed by FI×FI (53.1 ± 0.4) and DA×AR (55.0% ± 0.7). The colour of intra-region northern juveniles was higher than intra-region central juveniles (53.8% ± 0.2). Specifically, MA×MA was the darkest (53.9% ± 0.3), followed by WD×MA (53.5% ± 0.5). Colour of the inter-region juveniles was highest overall (54.2% ± 0.5). Specifically, DA×MA was the darkest (55.8% ± 0.8), followed by MA×DA (53.6% ± 0.6). There was no difference in colour between control FI×FI juveniles and DA×DA (P_e_ = 0.76, BGLMM).

#### d. Relationship between survival and growth

Relative to the native control juveniles (FI×FI), we observed a trade-off between growth and survival in the other juvenile crosses (Fig. 3). The control FI×FI juveniles exhibited the highest mean growth rate (248.6% ± 10.1 SE) but the second-lowest survival rates (29.0% ± 1.1). Also relative to FI×FI, all inter-region and northern intra-region crosses showed higher survival, but reduced growth rates. The magnitude of this trade-off differed by cross. Both the central (DA×DA) and northern (MA×MA) crosses had lower survival (DA×DA: 27.5% ± 0.9; MA×MA: 42.1% ± 1.0) and growth (DA×DA: 195.9% ± 8.0; MA×MA: 190.5% ± 6.1). In contrast, other intra-region crosses with parents from different reefs (WD×MA, DA×AR), showed increased survival relative to FI×FI (WD×MA: 59.5% ± 2.5; DA×AR: 69.3% ± 0.0), but 13.9% ± 17.6 lower growth (WD×MA: 234.7% ± 14.5; DA×AR: 207.9% ± 16.2). All inter-region crosses had higher survival compared to FI×FI but had reduced growth. In particular, MA×DA exhibited 29.6% ± 0.7 higher survival but 13.9% ± 17.9 lower growth, while DA×MA showed 49.2% ± 0.6 higher survival but 46.3% ±2 1.0 lower growth. Although the survival–growth relationship was not statistically significant (*p =* 0.79, linear regression), there was a subtle negative trend for the inter-region juveniles (WD×MA, MA×DA, DA×AR, DA×MA), but not intra-region juveniles (DA×DA, MA×MA).

**Figure 3:**
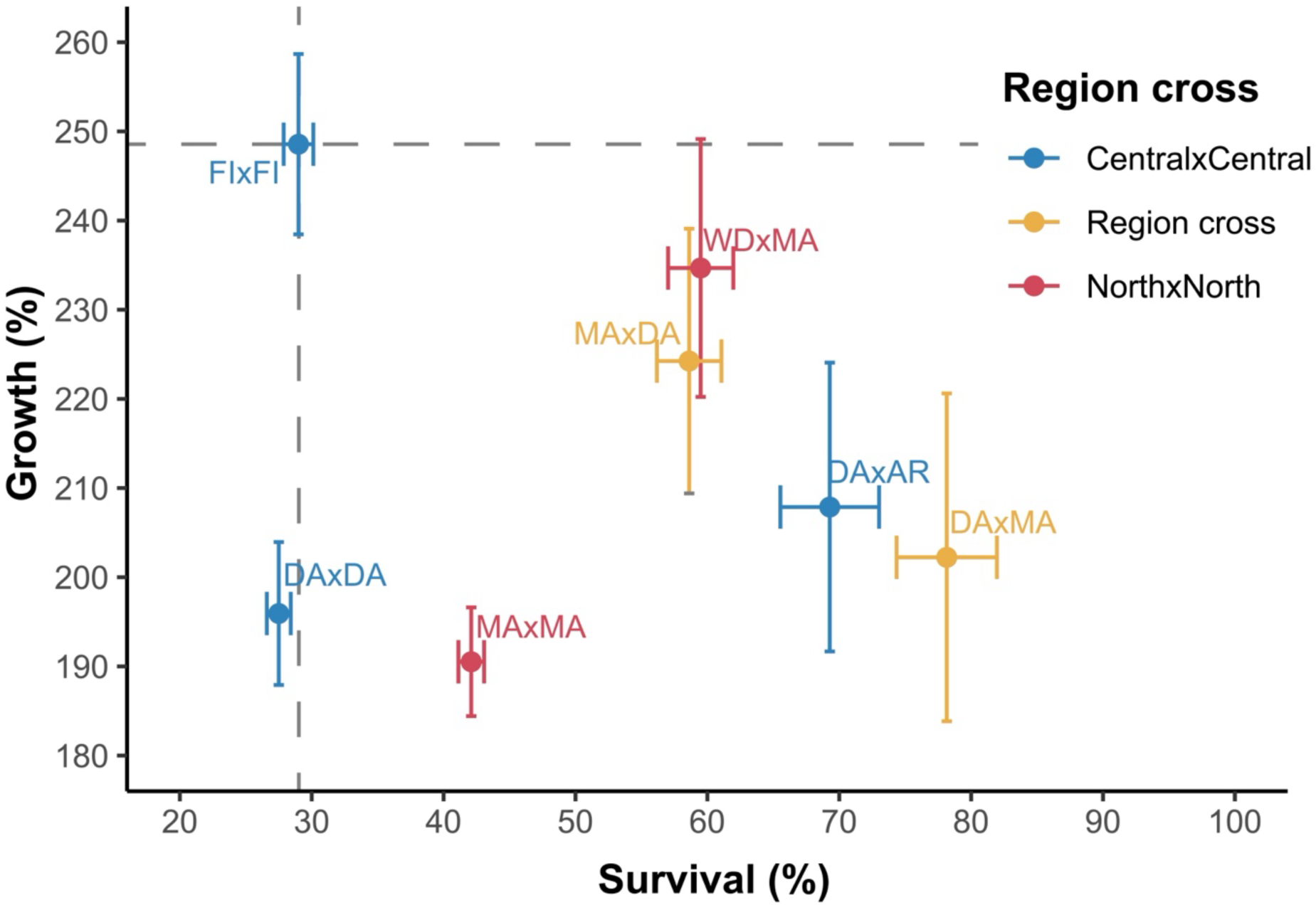
Relationship between survival (%) and growth (%) of outplanted *Acropora tersa* juveniles. Points represent the mean and error bars represent ± SE for growth rate and survival of juveniles in each reef cross over the deployment period and are coloured according to the regional cross. Grey dashed line correspond to the mean values from native control FI×FI juveniles (intra-region Fantome Island juveniles from the deployment site).

### Survival and bleaching of adult corals under experimental heat stress

Adult *A. tersa* colonies collected from northern reefs exhibited 3.75× higher survival odds at 32°C compared to adult corals collected from central reefs (Fig. 2D; Pₑ = 0.90, Bayesian Generalised Linear Mixed Model; BGLMM). Specifically, corals collected from Martin and Wood recorded the highest survival under heat stress (mean values for Martin = 96.7% ± 3.3 SE and Wood = 90.0% ± 5.6, respectively). The survival rates varied across the central reefs, with the highest survival of adult corals collected from Fantome (88.2% ± 5.6), followed by Arlington (75.0% ± 7.3). Adults from Davies (63.9% ± 8.1) recorded the lowest survival under heat stress conditions. All corals collected from all reefs recorded 100% ± 0 survival in the control treatment (27.5°C).

When comparing region-specific bleaching rates, northern *A. tersa* bleached less at 32°C compared to those from central reefs (Fig. 2E). At reef level, Fantome corals recorded the lowest bleaching and therefore highest colour scores (4.2 ± 0.1), followed by Martin (3.9 ± 0.1), Arlington (3.9 ± 0.1), Wood (3.5 ± 0.1), and Davies (3.4 ± 0.1; Supplementary Fig. S2D). When looking at change in colour, Martin bleached the least (Fig. 2E; −0.20 ± 0.1), followed by Fantome (−0.30 ± 0.1), and Wood (−0.50 ± 0.1). Corals from Davies bleached the most (−0.76 ± 0.1) followed by Arlington (−0.58 ± 0.1).

## DISCUSSION

Ocean warming is driving increasingly frequent and severe coral bleaching events (Frölicher et al., 2018; Oliver et al., 2018), threatening the persistence of coral reef ecosystems. Traditional conservation and restoration approaches alone are unlikely to secure the future of reefs under ongoing climate change (Anthony et al., 2017). In conjunction with strong and immediate emissions reductions, genetic interventions are being increasingly considered and may soon be necessary for reef management (Drury et al., 2022b). However, it is critical to rigorously assess both the risks and the benefits of such approaches, including AGF (Hamilton & Miller, 2016; Kovach et al., 2016), before considering widespread implementation. Here, we present the first field-based evidence that AGF can improve juvenile coral survival during a marine heatwave, providing essential empirical data for evaluating the feasibility of and implications for deploying genetic interventions on reefs.

### Assisted gene flow enhances juvenile survival during a marine heatwave

Juvenile corals deployed to Fantome Island, a reef on the central GBR that experienced >6 DHW of thermal stress, showed clear differences in survival amongst genetic crosses. Inter-region juveniles exhibited the highest survival, outperforming both northern and central intra-region juveniles. This result aligns with other laboratory studies which demonstrated enhanced tolerance of AGF offspring. This includes in *Platygyra daedalea* larvae crossed from the Arabian Gulf and Indian Ocean (Howells et al., 2021), intra-population breeding of heat tolerant Hawaiian *Montipora capitata* juveniles (but not in larvae; Drury et al., 2022a), *Acropora tersa*, *A. kenti*, *A. spathulata*, *A. millepora*, and *Goniastrea retiformis* larvae (Dixon et al., 2015; Macadam et al., 2025; Quigley et al., 2020a; Quigley et al., 2020b; Quigley & van Oppen, 2022; Weeriyanun et al., 2022), and in *Acropora tersa*, *A. kenti*, and *A. spathulata* juveniles (Macadam et al., 2025; Quigley et al., 2020b; Quigley & van Oppen, 2022) from the GBR. Notably, a study in the closely related *A. spathulata* also reported that some inter-region juveniles outperforming both parental region’s juveniles (Quigley et al., 2020b). The mechanism behind the greater survival of inter-region juveniles is unclear, but may be due to heterosis (hybrid vigour), which has been reported in in both animals (Cummins et al., 2025; Yang et al., 2024) and plants (Martins et al., 2019; Uwimana et al., 2012). Whether such benefits persist across generations or diminish is unclear (Hamilton & Miller, 2016; Kovach et al., 2016). Together, these findings position our study as the first to demonstrate enhanced survival of AGF corals directly on the reef under natural heatwave conditions, providing critical validation beyond controlled laboratory experiments and highlighting its value as a climate-adaptation strategy for reef restoration.

The increased survival demonstrated here provide novel and important field support for the efficacy of AGF during real marine heatwave events. Moreover, the magnitude of heat tolerance enhancement observed here exceeded those documented in the same coral species during laboratory heat-stress assays (Macadam et al., 2025). This suggests that laboratory experiments may be miscalculating the tolerance boost provided by selective breeding and likely reflecting factors available to wild corals but absent from laboratory experiments, including access to increased nutrients instead of filtered seawater, and additional diversity of symbiont communities rather than monocultures of laboratory-cultured symbionts. These conditions in the wild may explain the increased tolerance but require further work.

### Environmental drivers of local adaptation enhance heat tolerance

As expected, northern intra-region juveniles survived better compared to central intra-region juveniles, which aligns with expectations that corals from warmer reefs should bleach less (Hughes et al., 2017). This result also aligns with survival patterns of adult colonies from other studies (Liew et al., 2020; Putnam & Gates, 2015). It is important to note that substantial variation was also observed between reefs sourced from the same region, which again aligns with results from other laboratory studies both from within the GBR and other reef locations (Drury et al., 2022a; Humanes et al., 2022; Macadam et al., 2025; Marzonie et al., 2022; Naugle et al., 2024). Taken together, this suggests that the adult corals collected were likely locally adapted to their reef conditions.

We now know that the development of heat tolerance is shaped by a multitude of environmental factors, including mean monthly temperatures, daily temperature variation, and past history of temperature anomalies (Quigley & van Oppen, 2022). Moreover, other ecological and historical factors likely shape the development of heat tolerance (Naugle et al., 2024). Given that information, this could likely explain the differences in tolerances measured here among our central reef crosses. For example, the control cross – FI×FI – which was outplanted to its “home” reef, survived better compared to the other central cross (DA×DA). This suggests that the DA×DA juveniles, produced from the offshore central reef, may have been maladapted to Fantome’s inshore environment (characterized by higher temperature variability, turbidity, and nutrient loads) (Sully & van Woesik, 2020; Zweifler, 2024). This has also been noted in Florida reef environments (Kenkel et al., 2015). Interestingly, the higher survival and lower bleaching of Fantome adults did not align with the lower survival of the FI×FI juveniles. This suggests that ontogenetic differences, symbiont acquisition, or microhabitat variation may also be mediating the overall tolerance levels, which would be expected (Edmunds et al., 2004; Hazraty-Kari et al., 2022; Kenkel et al., 2015; McIlroy & Coffroth, 2017). Overall, these results highlight that likely broad regional environmental gradients as well as differences across life-history stages are important for determining the overall level of heat tolerance of individual corals.

### Potential for survival benefits to be counteracted by reduced growth under heat stress

Our results show the potential for a trade-off between enhanced survival but decreased growth. Although the overall trend was not statistically significant, there was a trend in, decreased survival with increased growth rate, specifically for the inter-region crosses but not the intra-region crosses. These results are important because relocating organisms into different habitats have the potential to generate fitness trade-offs, where gains in one trait come at the expense of another (Hereford, 2009). This is an area of active research in corals, where there is some evidence that some of the physiological mechanisms that enhance heat tolerance may carry metabolic costs that ultimately reduce growth or reproduction (Tomanek & Zuzow, 2010). Importantly, this trend wasn’t only measured in inter-region crosses, but it was most evident when examining the native FI×FI juveniles. These native juveniles exhibited the highest growth rates but comparatively lower survival during the heatwave. The other intra-region corals did not follow this trend, which instead showed low survival and low growth. In contrast, inter-region juveniles survived better but displayed reduced growth. Although this relationship was not statistically supported, the pattern aligns broadly with other findings in both adult (Cunning et al., 2015; Jones & Berkelmans, 2011; Roik et al., 2024) and juvenile corals (Strader & Quigley, 2022). Further work will be needed to understand under what conditions trait trade-offs may appear in corals, the mechanisms driving them, and the ecological implications.

The variation between crosses in survival and growth may also reflect differences in host and symbiont associations. It is well established that coral performance depends strongly on both the host genotype as well as by the algal symbiont community (Howells et al., 2013; Lin et al., 2019; Quigley et al., 2017; Sproles et al., 2020). Indeed, previous work has shown that the genetic background of the juvenile coral explains substantial variation in survival under heat stress, whereas symbiont assemblages explain variation in growth (Macadam et al., 2025; Quigley et al., 2020b). In this context, the increased growth of the native juveniles from FI×FI may reflect differences in symbiont communities or even patterns of local adaptation to the local symbiont communities available at Fantome Island. By extension, the non-native (northern and inter-region) juveniles may have experienced lower growth due to their exposure to “non-native” symbionts, leading to metabolic inefficiencies, symbiosis instability, or suboptimal nutrient exchange (Chan et al., 2024; Kenkel & Bay, 2018; Lin et al., 2019; Morris et al., 2019; Sproles et al., 2020). Differences in heterotrophy or symbiont composition between native and non-native corals may further contribute to these patterns (Anthony & Fabricius, 2000; Fox et al., 2018; Jones & Berkelmans, 2010; Jones & Berkelmans, 2011). Importantly, although AGF corals here exhibited enhanced survival during a heatwave, their slower growth could limit recovery, a well-recognized ecological challenge for reefs (Álvarez-Noriega et al., 2023) and underscores the need to better understand how genetics and the environment interact to shape coral physiology under stress and long-term coral reef persistence.

### Implications for coral reef restoration

Although survival rates for one-year-old juvenile corals are challenging to estimate, it is likely low (0.6–2%; Doropoulos et al., 2019) and non-linear (Cameron & Harrison, 2020). Interestingly, the survival measured in this study exceeded some past estimates (Cameron & Harrison, 2020) but was comparable to other field trails (Quigley et al., 2021). The higher survival could in part be due to the use of predator-deterring devices to shield the coral juveniles (Whitman et al., 2024). These results underscore that, alongside thermal history of the adult corals used in breeding, the ecological interactions should be accounted for when evaluating AGF outcomes.

Aside from accounting for other ecological interactions like predation, our findings also highlight several key considerations for potential future deployment of AGF corals as a conservation tool. First, AGF success will depend on accurately forecasting thermal conditions at recipient sites (Aitken & Whitlock, 2013). Mismatches between expected and actual conditions could lead to maladaptation (Torda & Quigley, 2022). For example, we found increased heat tolerance accompanied by varying degrees of reduced growth. Second, while some variation in underlying genetics is necessary to introduce novel genetic variation to populations (Hamilton & Miller, 2016), excessive genetic divergence may disrupt locally adapted traits or co-adapted gene complexes (Kovach et al., 2016). Not only did we see evidence of local adaptation here, but recent genetic evidence also suggests that fine-scale genetic structure does exist in at least some coral species across the GBR (Cooke et al., 2020; Matias et al., 2023). Finally, we detected no evidence of incompatibility, in contrast to findings in other marine taxa (Cummins et al., 2025; Pregler et al., 2023). However, further work is needed to define species boundaries and quantify genetic differentiation, which will be critical for optimizing the use of conservation tools like AGF. In summary, this study provides robust field-based evidence that AGF can enhance coral thermal tolerance during marine heatwaves in the wild. By combining insights of local reef environments and trait performance our results offer important insights into how AGF corals might perform in the wild under future climate warming scenarios.

## DATA ACCESSIBILITY STATEMENT

All data will be available on https://github.com/alexmacadam241/AGF-Outplanting.

## AUTHOR CONTRIBUTIONS

**Alex Macadam:** Conceptualization, Methodology, Formal analysis, Investigation, Data curation, Writing – original draft, Writing – review & editing, Visualization, Project administration. **Carys Morgans:** Investigation. **Ira Cooke:** Validation, Writing – review and editing, Supervision. **Patrick Laffy:** Validation, Writing – review and editing, Supervision. **Jan Strugnell:** Validation, Writing – review and editing, Supervision. **Andrea Severati:** Conceptualization, Methodology, Resources, Project administration. **Kate Quigley:** Conceptualization, Methodology, Investigation, Validation, Resources, Writing – original draft, Writing – review & editing, Supervision, Project administration, Funding acquisition.

## ACKNOWLEDGMENTS

We acknowledge the Wuthathi, Gunggandji, Wulgurukaba, and the people of the Hopevale Congress Aboriginal Corporation and Palm Island Improved Land Management Practices ILUA as the Traditional Custodians of sea Country where this research took place. The authors acknowledge their Elders past, present, and emerging, and their continuing spiritual connection to sea Country. Coral colonies were collected under permit G22/46111.1 issued to K.M.Q by the Great Barrier Reef Marine Park Authority. This research was funded by the Australian Institute of Marine Science, the Great Barrier Reef Foundation and the Reef Restoration and Adaptation Programme, a partnership between the Australian Government’s Reef Trust and the Great Barrier Reef Foundation. We thank Sam Noonan, Taylor Whitman, Maren Toor, Carlos Alvarez Roa, Saskia Jurriaans, Rebecca Forster, and RV Cape Ferguson crew for field assistance.

## SUPPLEMENTARY MATERIAL

**Supplementary Figure S1:**
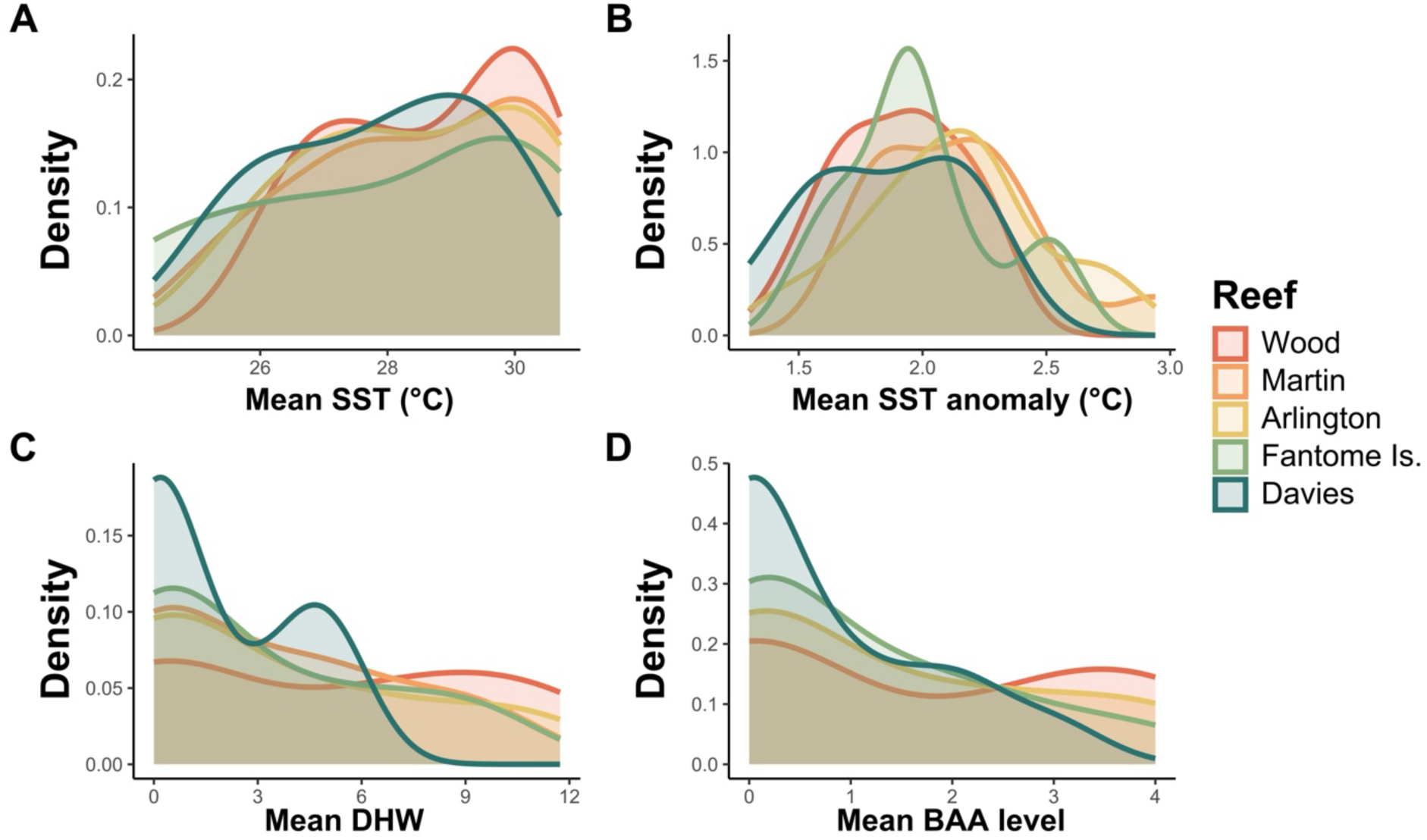
Historic temperatures of collection sites from 2016 - 2021. (A) Density of mean historic sea surface temperatures (SST; 2016 - 2021) for each collection reef. (B) Density of mean historic SST anomaly (2016 - 2021) for each collection reef. (C) Density of mean degree heating weeks (DHW; 2016 - 2021) for each collection reef. (D) Mean Bleaching Alert Area (BAA) for each collection reef. Temperature measurement resolution was one measurement every 3 days.

**Supplementary Figure S2:**
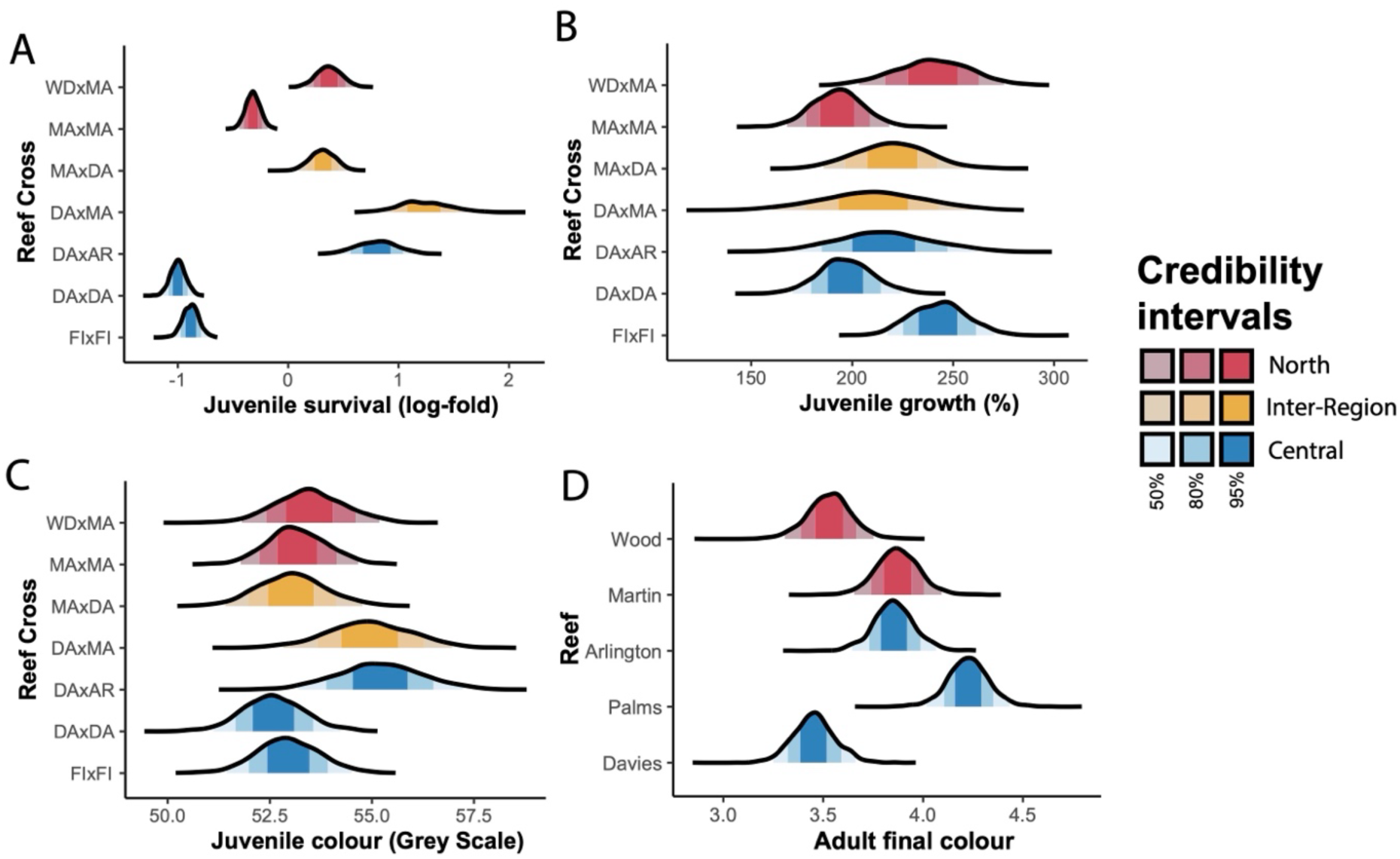
Distribution of estimated marginal means (EMM) of parameters of *Acropora tersa* adult and juvenile corals. EMM for: (A) survival (log-odds) of juvenile corals at the end of deployment (95 days) by reef cross, (B) growth (%) of juvenile corals at the end of deployment (95 days). (C) final colour (grey scale as a proxy for bleaching score) of juvenile corals at the end of deployment, (D) final colour (Coral Health Chart colour score D scale) of adult corals in the hot (32°C) treatment. Higher grey scale percent means more pigmentation. Colours represent the region; red = north, blue = central, and yellow = inter-region crosses. Intervals represent EMM 50%, 80% and 95% credibility intervals.

**Supplementary Table 1:**
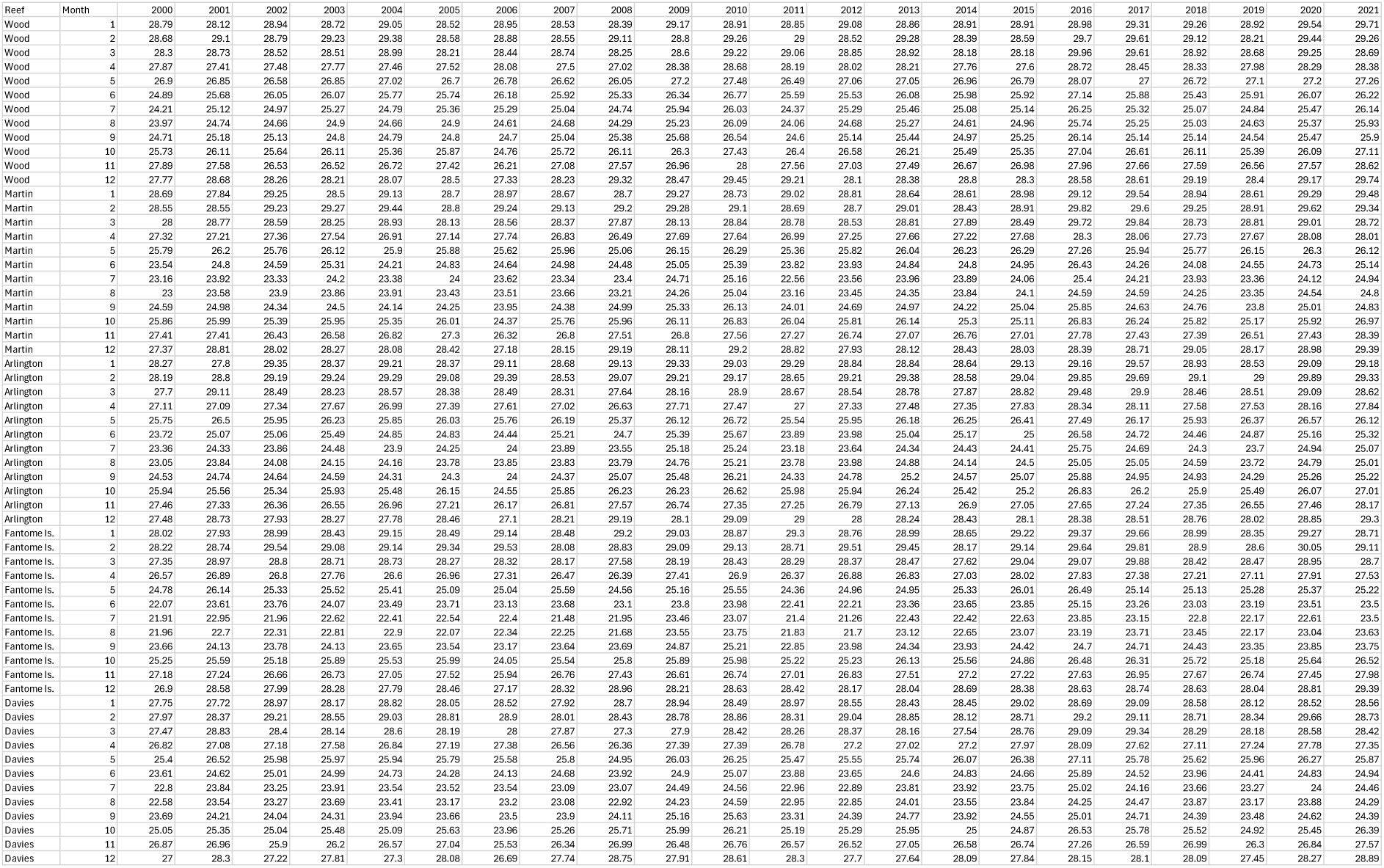
Monthly maximum mean for each collection site.

**Supplementary Table 2:**
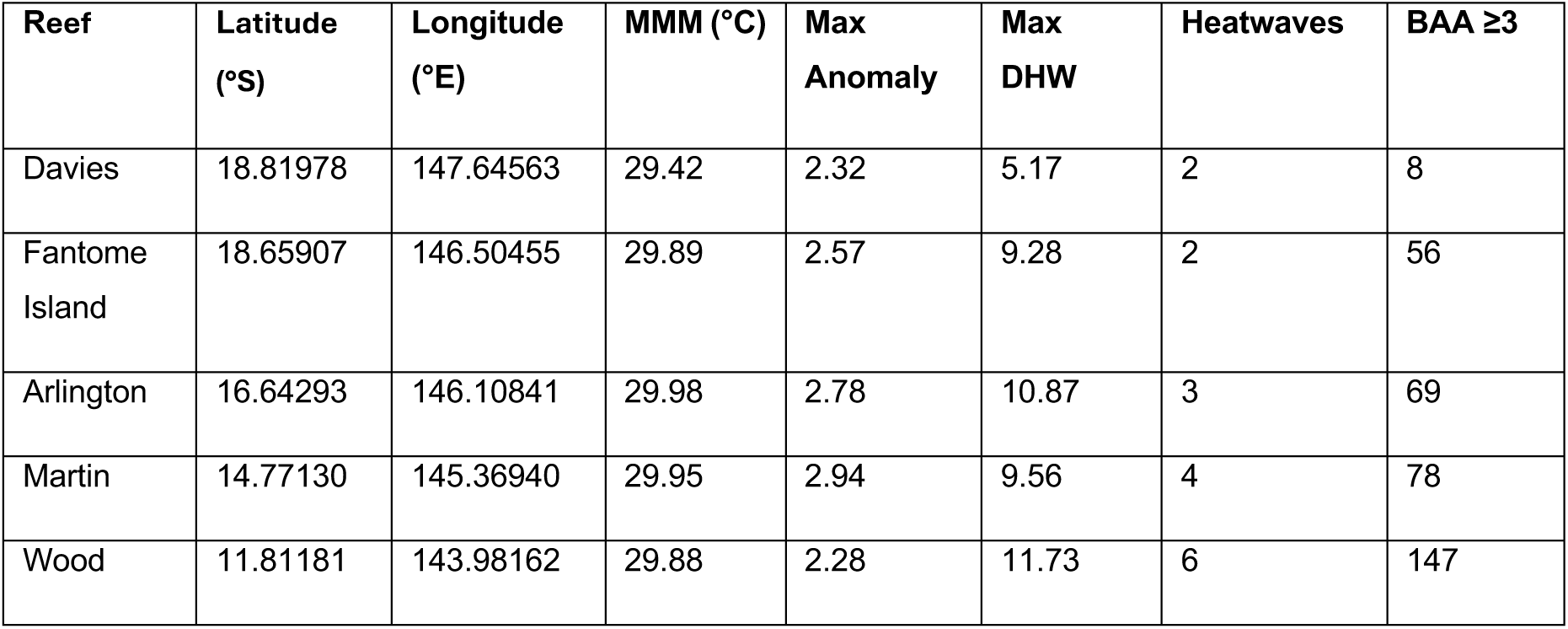
Locations and environmental conditions for collection sites of gravid coral colonies used for Assisted Gene Flow experiments in the Great Barrier Reef Marine Park (GBRMP) using NOAA Coral Reef Watch data. Here, maximum monthly mean temperature (MMM, °C) is defined as the average SST of the hottest month in each year between 2000 and 2021. Maximum anomaly is the maximum sea surface temperature anomaly encountered between 2000 and 2021. An anomaly is defined as the difference between observed temperature and the long-term average for that day. Maximum degree heating weeks (DHW) is the maximum DHW experienced between 2000 and 2021. DHW shows accumulation of heat stress intensity and duration. A DHW is defined as one week that SST is 1°C above the local highest summertime mean. Heatwaves records the number of heatwaves (defined as occurrences of at least 4 DHWs within a 12-week period) between 2000 and 2021. Bleaching alert area (BAA) >2 shows the number of days spent at or above level 3 (Bleaching alert level 1: SST above bleaching threshold, DHW ≥4) between 2016 and 2021.

